# Development and characterization of phytosterol nanoemulsions and self-microemulsifying drug delivery systems

**DOI:** 10.1101/585166

**Authors:** Yuan Chuanxun, Zhang Xueru, Jin Risheng

## Abstract

The aim of this study is to develop a self microemulsion drug delivery system for phytosterols to improve the solubility and bioavailability. The results showed that the formulation of phytosterol self-microemulsion is: lemon essential oil in oil phase, polyoxyethylene hydrogenated castor oil 40 and Tween 60 in emulsifier, polyethylene glycol 400 in co-emulsifier, Km = 7:3, Kp = 3:1, Ke = 50%. The drug loading of phytosterol self-microemulsion prepared by this method was 87.22 mg/g, encapsulation efficiency was 89.65%, particle size was 48.85nm, potential was −12.863mV. In vitro release experiment showed that the release of phytosterols in microemulsion was more than 90%, and the release curve was in accordance with the first-order kinetics equation. The pharmacokinetic analysis of PSSM synthesized by this method shows that PSSM can increase the bioavailability of PS more than three times, so it is necessary to do more in-depth research on the self-microemulsion delivery system of phytosterols.

## Introduction

Phytosterols(PS) are triterpenes, an important component of plant cells, and are commonly found in seeds, vegetable oils and grains. Common in nature are β-sitosterol, campesterol, stigmasterol and ergosterol [1]. It has the physiological functions of lowering blood lipid [2, 3], lowering cholesterol [4–6], preventing cardiovascular disease [7],anti-cancer[8], anti-oxidation, scavenging free radicals[9, 10], enhancing immunity[11], anti-inflammatory [12, 13] and so on. Especially in reducing blood lipids, PS and cholesterol have similar molecular structure, which can competitively inhibit the absorption of cholesterol in the human intestinal tract, thereby reducing blood lipids [14]. Studies have shown that taking 2 g of PS a day can reduce low density lipoprotein(LDL)by 8.8 %[15].Like unsaturated fatty acids and cholesterol, PS is easy to oxidize and can produce various phytosterol oxidation products (POPs), such as hydroxyl groups, epoxy resins, ketones and triol derivatives, especially when heated or stored for a long time[16].PSs are insoluble in water and slightly soluble in oil, which limits their application in the pharmaceutical and health products industries. Self-microemulsion drug delivery system (SMEDDS) improves the bioavailability of drugs by improving their solubility, lymphatic absorption and intestinal permeability[17].In addition, the microemulsion has good stability[18], which can reduce the oxidation deterioration caused by the storage of PS. In this study, phytosterol self-microemulsion (PSSM) was prepared, the release characteristics of phytosterol in vitro and the pharmacokinetic analysis were investigated by reverse dialysis, which provided a theoretical basi for the study of PSSM.

## Materials and methods

### Materials

Phytosterol with purity of 98% was purchased from Xi’an Ruiying Biotechnology Co., Ltd. (Xi’an, China). Polyoxyethylene hydrogenated castor oil 40(HCO-40), tween 20, tween 60, tween 80, isopropyl myristate(IPM), 1,2-propanediol, polyethylene glycol 400 (PEG400) and polyoxyethylene castor oil (EL) were obtained from Shandong Yousuo Chemical Technology Co., Ltd. (Shandong, China). Lemon essential oil(LEO), linoleic acid(LA), oleic acid(OA) and ethyl oleate(EO) were purchased from Shandong Baihong New Materials Co., Ltd. Methanol (chromatographic purity, Tedia Company); the other reagents are analytical reagents. The experimental water is distilled water.

### Oil-water partition coefficiency

10 ml n-octanol was measured and placed in three conical bottles. Distilled water, 0.1 mol/lHCL and pH6.8 phosphate buffer(pH6.8 PBS) were added to each bottle. The two phases were saturated and separated overnight. The saturated n-octanol phase and three medium water phases were obtained[19]. The drug was dissolved in n-octanol until saturated.The supernatant was centrifuged (5000r/min, 10min) and the original solution was obtained. 2 ml of original solution and 2ml of the above three saturated water were mixed and kept for the night.The drug concentrations in original solution and three saturated water phases were determined by HPLC. The formula of oil-water partition coefficient (P) was as follows:

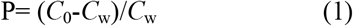

where C_0_ represents the drug concentration in original solution; C_w_ is the drug concentration in the aqueous phase after equilibrium.

### Solubility studies

PS was added to different oil phases and emulsifiers to saturate, shaking for 48 h (37 °C 100 r / min), static equilibrium for 24 h, centrifugation (5000 r / min, 10 min). The upper liquid was diluted 10 times and detected by HPLC.

### Pseudo-ternary Phase Diagram

11 phase and emulsifier with high solubility were selected to combine with co-emulsifier. The self-microemulsion system was formed by mixing the oil phase with its mass ratio of 9:1, 8:2, 7:3, 6:4, 5:5, 4:6, 3:7, 2:8, 1:9.With water drop method, visual observation and laser pen assistance, the critical point of clarification and path formation was recorded. The percentage of oil phase, emulsifier phase and water phase in the whole system was calculated, and pseudo-ternary phase diagram was drawn[20].

### Preparation of PSSM

Adding 4.00g LEO, 2.40g HCO-40, 1.03g Tween 60, 1.14g PEG400 and 2.00g PS, shaking in shaking flask (37 C, 100r/min) for 48 hours, standing for 24 hours, removing undissolved PS, and obtaining PSSM[21].

### Particle size and zeta-potential

The prepared SM was diluted 50 times with distilled water at 37°C. The particle size, polydispersity index (PDI) and Zeta potential of the microemulsion were measured. All samples were balanced for 120s at 25 °C before measurement and each sample was measured three times in parallel with Zetasizer Nano S90 High Sensitivity Nano Particle Size Analyzer (UK Malvern Instruments Co., Ltd.)[22].

### Drug loading and entrapment efficiency

SM 25.00 mg in 10 ml volume bottle was shaken with methanol until calibration, ultrasound was performed for 10 minutes, concentration was determined by HPLC and recorded as W1[23]. 100.00 mg SM in 5 ml volume bottle was shaken with distilled water to scale. After 12 hours of dialysis(MWCO 500 Da), the SM was diluted with methanol for 10 times and demulsified by ultrasound for 30 minutes then W2 was determined by HPLC. The formula for determination of drug loading(DL) and entrapment efficiency (EE) are as follow[24]:

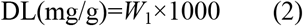

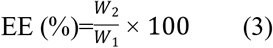

where *W*_1_ is PS content in self microemulsion, and *W*_2_ is PS content in microemulsion.

### Morphology observation

The morphology of PSSM prepared under optimum conditions was observed by transmission electron microscopy (TEM). Test conditions: 200 kV acceleration voltage, No. 1 condenser aperture.Take 0.1g SM, dilute it 50 times with distilled water at 37° C, put a drop on copper, absorb excess liquid with filter paper, dye it with 2% phosphotungstic acid solution and dry it, and observe it under JEM-2100F Field Emission Transmission Electron Microscope (Japan)[12].

### Stability of PSSM

Three samples of the same PSSM were stored at 4, 25 and 40 degrees for 45 days. The particle size changes were detected every 5 days and the precipitation degree of the drug during storage was observed.

### In vitro release study

Study on in vitro release of PSSM and phytosterol raw materials(PSRM) by reverse dialysis, the dilution media were distilled water, pH 6.8PBS and 0.1 mol/L HCL solution, the parameters of magnetic stirring water bath were 37 ° C, 50 r/min. Ten dialysis bags (MWCO 12,000-14,000 Da) were added with 5 ml dilution medium, balanced for 12 h, 4.00 g SM in 900 ml dilution medium with 1% Tween 80. Take out a dialysis bag respectively at appropriate intervals and 5 ml of the same release medium was supplemented[25]. The drug concentration in dialysis bags was determined by HPLC, and the drug release curve was drawn.

### Pharmacokinetic study

Twelve male SPF Sprague-Dawley rats (240+10 g) were fed adaptively for 7 days. The temperature and humidity of the animal room were 25±2°C, 55±5% and 12/12 h light-dark cycles. All animals used in the study were treated in strict accordance with the laboratory animal care and use guidelines of the National Institutes of Health. All procedures were approved by the Animal Nursing Assessment Committee of Hefei University of Technology, China.

SD rats were randomly divided into PSSM and PSRM groups, two groups were fed with the same amount of PS. Six rats in each group were fed 10 mg/g (PS content is 87.22 mg/g), fasting 24 hours before administration and drinking water freely. At the appropriate time point, about 0.3 mL of blood was collected from the posterior orbital venous plexus of rats and placed in a heparinized centrifugal tube. The plasma was separated by centrifugation at 6000 rpm for 10 minutes and stored at −80 °C[26]. The content of PS in plasma was determined by HPLC, and the data of plasma concentration were processed by WinNonlin 5.2 software[27]. The pharmacokinetic parameters are: area under curve (AUC); peak concentration (Cmax); time peak concentration (tmax) and relative bioavailability (F%). The formulas are as follows [28, 29]:

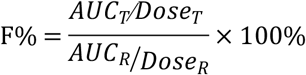

where *T* refers to the test preparation, *R* is the reference preparation.

### Conditions of HPLC

Chromatographic column: Eclipse plus C18 (Agilent, 4.6 × 250mm/5μm); mobile phase: methanol: water = 99:1; flow rate: 1.0mL/min; detection wavelength: 210 nm; column temperature: 30°C; injection volume: 10μL.

## Results and discussion

### Oil-water partition coefficiency

The drug distribution coefficient P in n-octanol-water system is a physical constant that simulates the lipophilicity or hydrophobicity of drugs in vivo, and is used to predict the absorption of drugs in the intestine[30]. When 0 < logP < 3, the drug can be absorbed by gastrointestinal tract, when logP < 0, the drug is hydrophilic; when logP > 3, the drug is lipophilic, but neither of them is easily absorbed by gastrointestinal tract.It is generally believed that logP between 2 and 3 is easily absorbed, while drugs with logP < 2 are not easily absorbed[31, 32]. Table 1 shows that the logP of phytosterols in the three media is less than 2, indicating that phytosterols are difficult to be absorbed by intestine. Therefore, it is necessary to improve the dosage forms of phytosterols.

**Table 1.** Apparent partition coefficient of phytosterol in different solvents

| solvent | distilled water | 0.1mol/L HCL | pH6.8PBS |
| --- | --- | --- | --- |
| *P* | 1.762 | 41.69 | 12.88 |
| Log*P* | 0.246 | 1.62 | 1.11 |

### Solubility studies

The oil phase selected by SMEDDS generally has high solubility for drugs[33]. The solubilities of LEO, LA, EO, OA and IPM were determined. As shown in Fig. 1, the solubilities of different oils to phytosterols were as follows: LEO > OA > IPM > LA > EO.Surfactants have certain toxicity. Generally, polyoxyethylene vegetable oil is less toxic than single-chain surfactants and lipid surfactants are less toxic than ethers. Non-ionic surfactants are preferred for oral SM.Commonly used nonionic surfactants are tween, polyoxyethylene, polyethylene glycerol, Sipan and natural lecithin[34–36].In this experiment, Tween 20, Tween 60, Tween 80, HCO-40 and EL were selected to determine the solubility of PS. The results showed that Tween 60 > Tween 80 > polyoxyethylene castor oil > polyoxyethylene hydrogenated castor oil > Tween 20 (Fig. 2). So LEO and tween 60 were selected as the oil phase and emulsifier of PSSM. After many preliminary experiments, it was found that the particle size of the prepared microemulsion was larger than 300 nm.But after adding HCO-40 in microemulsion, the particle size of microemulsion decreases obviously. The reason may be that HCO-40 and Tween 60 have complementary hydrophilic group conformations and play a synergistic role in compound emulsifiers (this part may be further discussed in future research). In summary, the PSSM oil phase is lemon essential oil, and the emulsifiers are compounds of HCO-40 and Tween 60.

**Fig 1.**
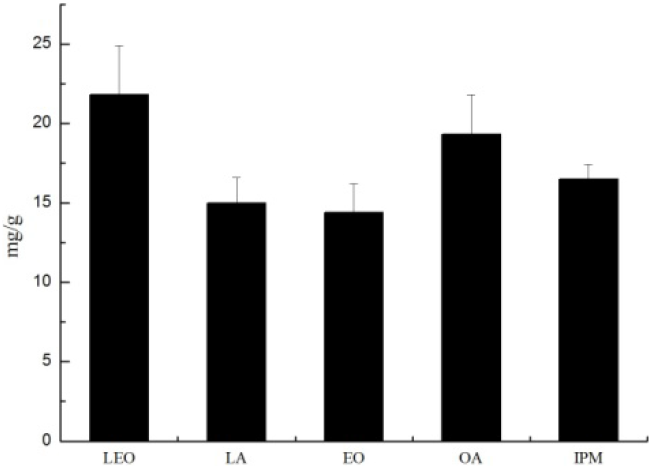
Solubility of phytosterols in different oil phases

**Fig 2.**
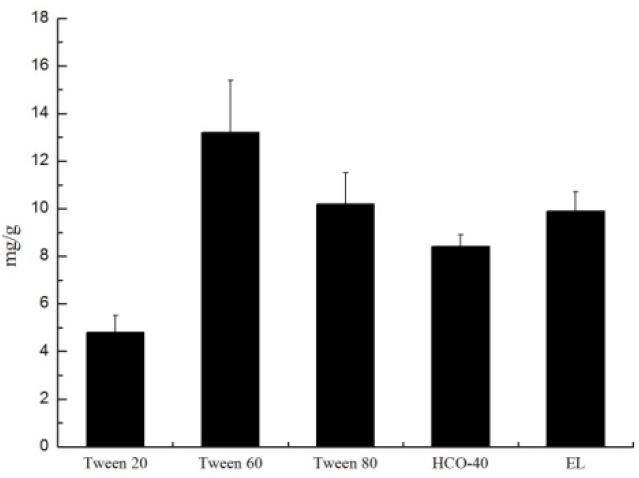
Solubility of phytosterols in different emulsifiers

### Pseudo-ternary phase diagram

The larger the area of microemulsion in pseudo-ternary phase diagrams, the easier the system to form microemulsion and the more stable it is[37, 38].In this part of the experiment, the single factor variable method was used to select the appropriate co-emulsifier and its content with the area of microemulsion as the index.2.00g LEO, 1.50g HCO-40 and 1.50g Tween 60 were added with 1.00g absolute ethanol, 1,2-propanediol and PEG400 to determine the appropriate co-emulsifier, or 12.00g, 9.00g, 6.00g, 3.00g, 1.50g, 1.00g, 0.75g co-emulsifier, that is, the ratio of emulsifier to co-emulsifier (Kp) = 1:4, 1:3, 1:1, 2:1, 3:1, 4:1, respectively, to determine the appropriate Kp.As can be seen from Figs. 3 and Table 2, when PEG400 is used as co-emulsifier and Kp=3:1, the proportion of microemulsion area in pseudo-ternary phase diagram is the largest. Therefore, PEG400 was chosen as emulsifier and Kp=3:1.

**Fig 3.**
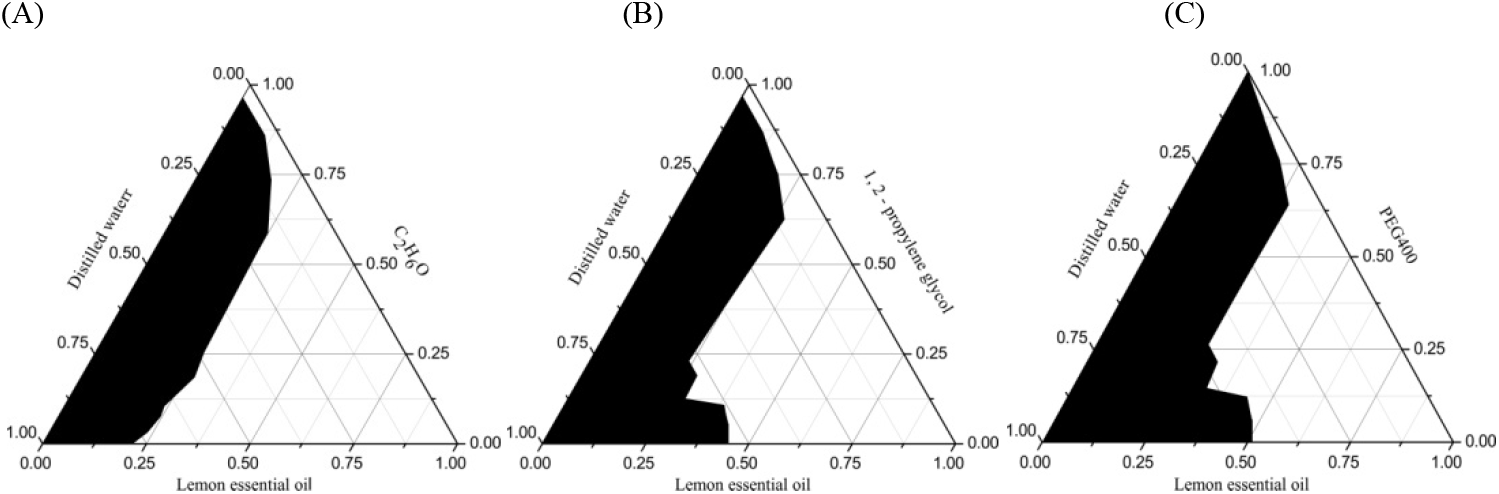
Pseudoternary phase diagrams were drawn with different co-emulsifiers. (A) Anhydrous ethanol; (B) 1, 2 - propylene glycol;(C) PEG400.(The shaded part of the pseudo-ternary phase diagram is the microemulsion region)

**Table 2.** The proportion of microemulsionb in pseudo ternary phase diagrams with different Km

| $K_p$ | 1:4 | 1:3 | 1:2 | 1:1 | 2:1 | 3:1 | 4:1 |
| --- | --- | --- | --- | --- | --- | --- | --- |
| Microemulsion | 49.31% | 55.72% | 53.88% | 51.12% | 56.04% | 58.24% | 51.97% |

Table 2 are the percentages of shadow area in the corresponding pseudo-ternary phase diagram calculated by AutoCAD 2012 software in the total pseudo-ternary phase diagram area.

### Particle size and zeta-potential

SMEDDS is a thermodynamic and dynamic stabilization system which can form particle size 10-100nm after emulsification. Particle size distribution is a standard for measuring the formation of microemulsion[39, 40]. Polydispersity index (PDI) is an indicator of particle size distribution. It should generally be between 0 and 0.3, and should not exceed 0.7. The smaller the value, the more concentrated the particle size distribution and the more uniform.Zeta potential is one of the indexes to determine the physical stability of microemulsion, and it is an index to measure the intensity of mutual exclusion or attraction between particles. The greater the zeta potential, the more stable the system is. The particle size of microemulsion is related to the speed and degree of drug release. The smaller the particle size, the larger the interface area, and the easier the diffusion to gastrointestinal juice, the more complete the drug release[41].

2.00g LEO, 1.00G PEG400, 3.00g emulsifiers were used, in which the mass ratio of HCO-40 to Tween60 (Km) was 1:9, 2:8, 3:7, 4:6, 5:5, 6:4, 7:3, 8:2, 9:1, diluted 50 times with distilled water. The optimum Km was determined by single factor variable method with particle size, PDI and zeta potential as indexes. Similarly, take 2.00 g LEO and 1.00 g PEG400, respectively, and take the emulsifier to the total system mass ratio (Ke) of 20%, 30%, 40%, 50%, 60%, 70%, 80%, Km = 5:5, to study the best Ke. From Fig. 4 and 5, the best Km = 7:3 and the best Ke = 50%.In conclusion, the formulation of PSSM is: LEO in oil phase, HCO-40 and Tween 60 in emulsifier, PEG400 in co-emulsifier, Km = 7:3, Kp = 3:1, Ke = 50%.

**Fig 4.**
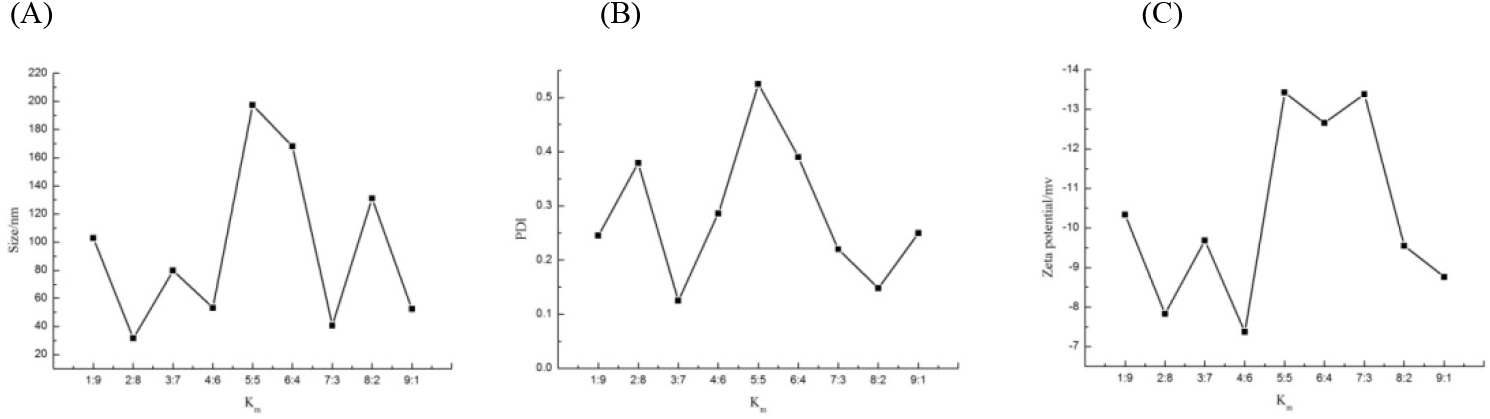
The relation between Km and particle size, PDI, potential. (A) particle size; (B) PDI; (C) potential.

**Fig 5.**
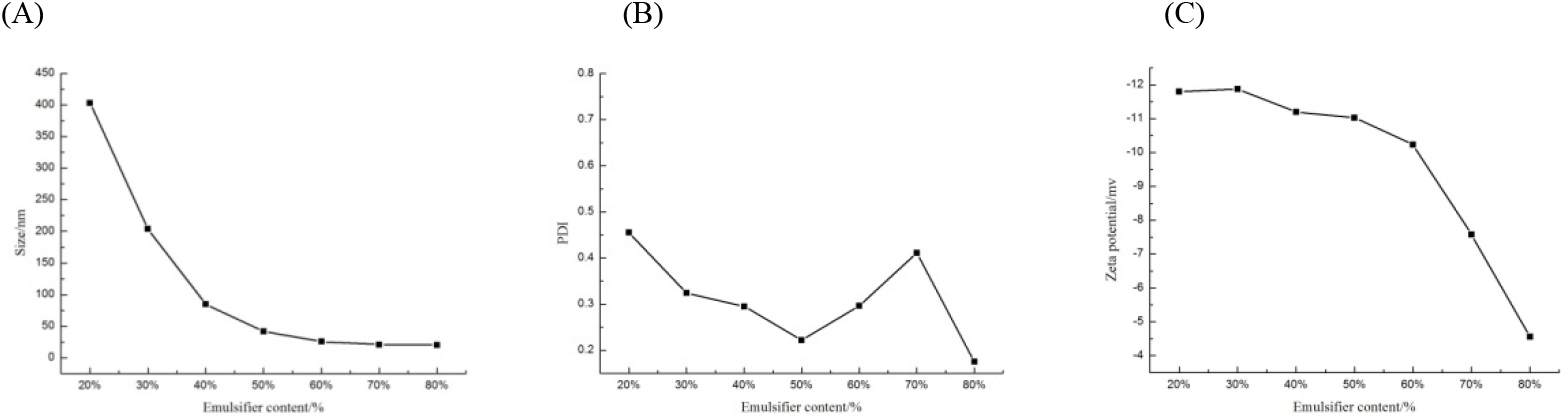
The relation between Ke and particle size, PDI, potential. (A) particle size; (B) PDI; (C) potential.

### Evaluation of PSSM

#### Drug loading and encapsulation rate

Drug loading refers to the ratio of the drug content in the system to the total weight of the system. Encapsulation ratio refers to the ratio of the total drug quantity encapsulated in the system to the total drug input in the system[42]. In this part, the drug loading, encapsulation efficiency, particle size, PDI and potential of the prepared PSSM were tested. The results were shown in table 3, and the particle size and potential distribution were shown in Fig. 6 and 7.

**Table 3.** The drug loading, encapsulation efficiency, particle size, PDI and potential of the PSSM

| DL(mg/g) | EE (%) | Particle size (nm) | PDI | Potential (mv) |
| --- | --- | --- | --- | --- |
| 87.22 | 89.65 | 48.85 | 0.293 | -12.863 |

**Fig 6.**
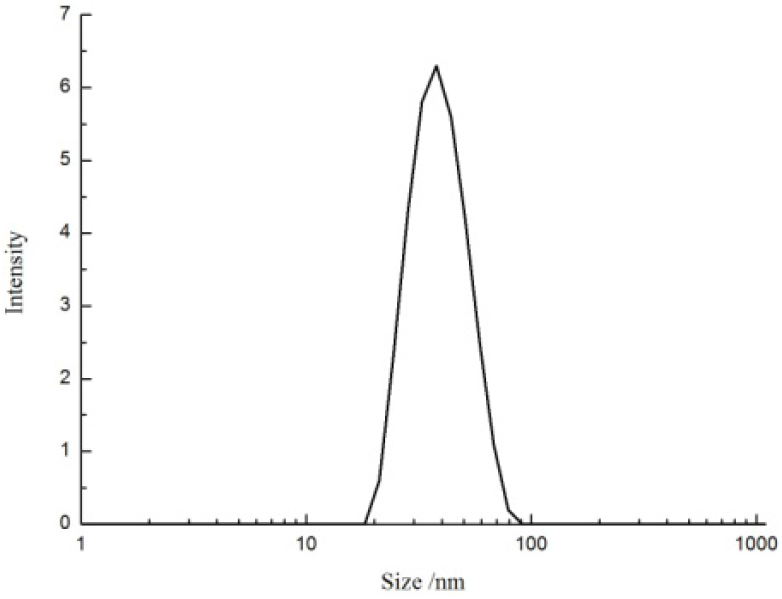
Particle size distribution

**Fig 7.**
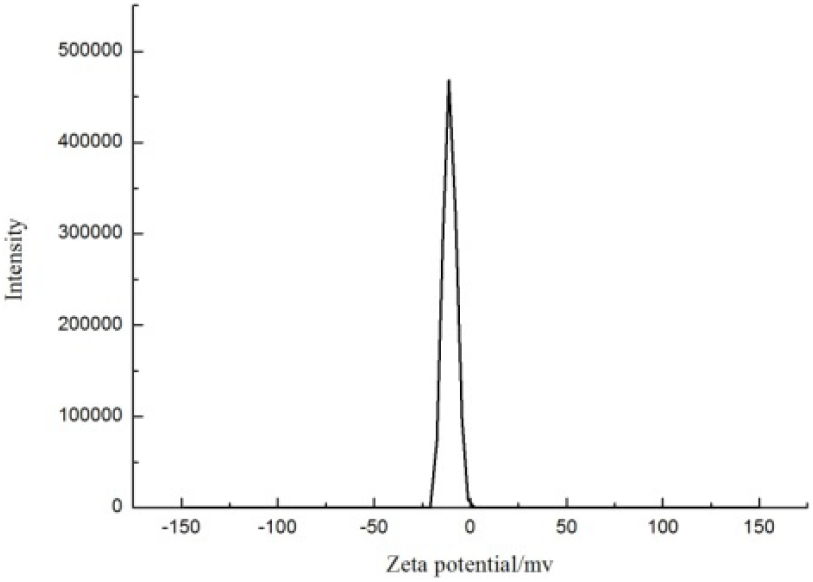
Zeta potential distribution diagram

#### Morphology observation

The PSSM was diluted 50 times with distilled water and observed by TEM. In Fig. 8(A) and (B) are microemulsions with different magnifications enlarged under the same field of vision. From Fig. 8(A), we can see that the particle size of microemulsions is relatively uniform and evenly distributed in the whole field of vision. as can be seen from Fig. 8(B), the morphology of the microemulsion is spherical, and there is little adhesion between microemulsion particles. The particle size is about 48 nm, which is consistent with the particle size measured by Zeta potentiometer.

**Fig 8.**
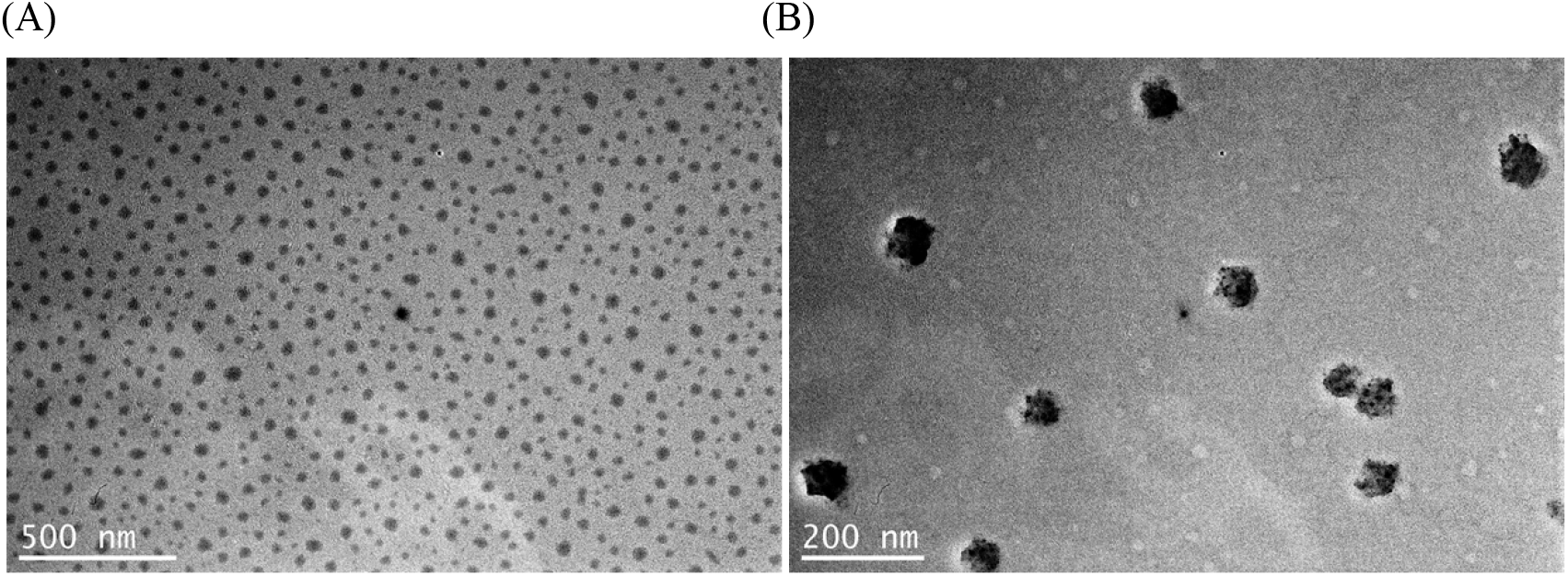
TEM micrograph of the PSSM. (A) TEM micrograph of the PSSM at magnitude 20,000×. (B) TEM micrograph of the PSSM at magnitude 50,000×.

#### Stability of PSSM

The particle size change of PSSM during storage in different environments is shown in Fig. 9. It can be seen that the particle size of PSSM varies significantly at 4° C and 40° C than at 25° C, the particle size change range is relatively small. As can be seen from Fig. 10, PSSM did not precipitate in different environments for 45 days. In conclusion, PSSM has good storage stability.

**Fig 9.**
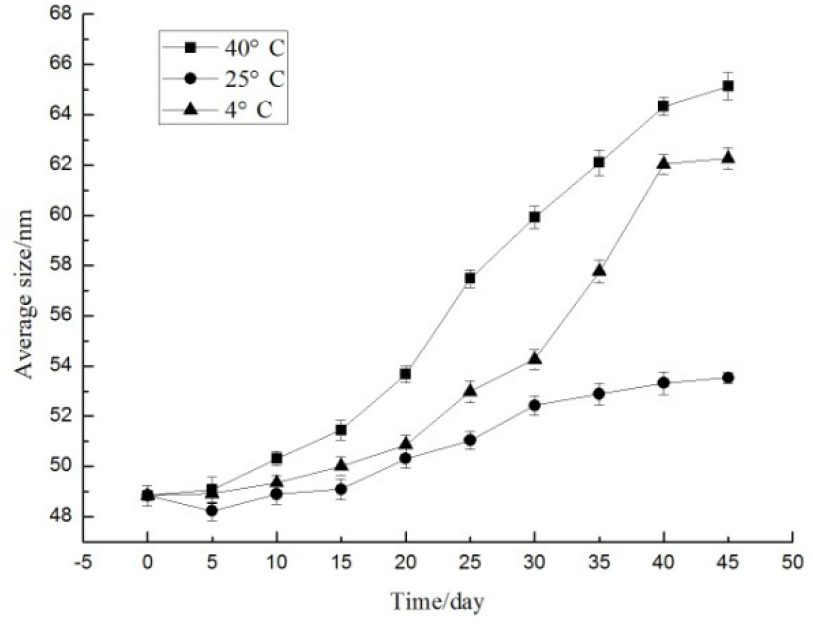
Particle size change during storage of PSSM

**Fig 10.**
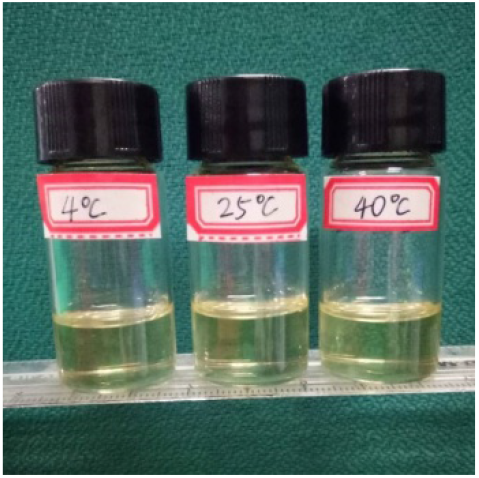
PSSM after 45 days storage

#### In vitro release study

In vitro drug release is an important index for evaluating the drug delivery system of granules. The drug release behavior and mechanism in vivo can be predicted by in vitro drug release experiments, and the structure of granules can be understood. At present, dialysis, reverse dialysis, ultrafiltration and Franz diffusion cell methods are commonly used to evaluate the in vitro release of SMEDDS[43, 44]. The similarity of release curves was compared by similarity factor (f2) method with DDSolver software and the drug release equation was fitted[45]. Fig. 11 shows that PSSM release completely within 3h in three dilution media and there is no significant difference in drug release. Table 4 shows that 50 < F2 < 100, indicating the three release curves are similar to each other [46]. Fig. 12 shows that PSRM almost completely released in about an hour, and the release rate was higher than that of PSSM.The results show that the release mechanism of PSSM conforms to the first-order kinetics equation (Table 5).

**Fig 11.**
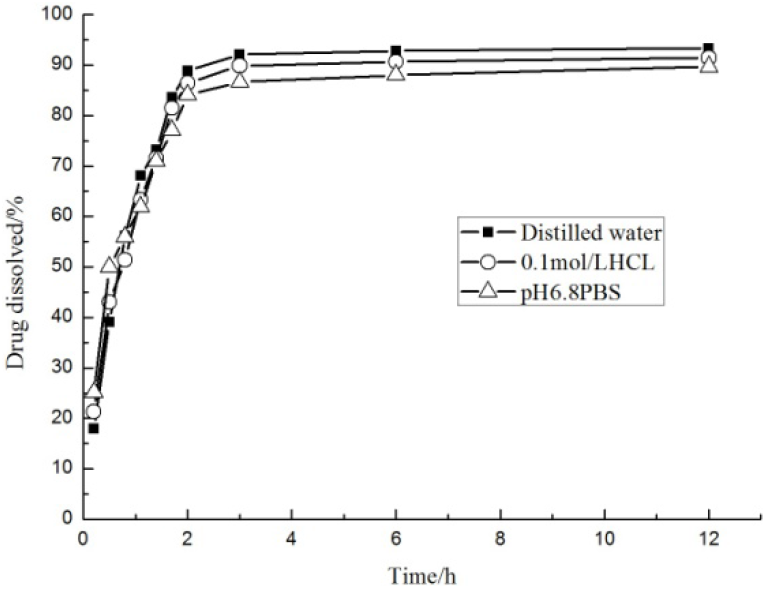
Release curve of PSSM in different dilution media pH 6.8 PBS

**Fig 12.**
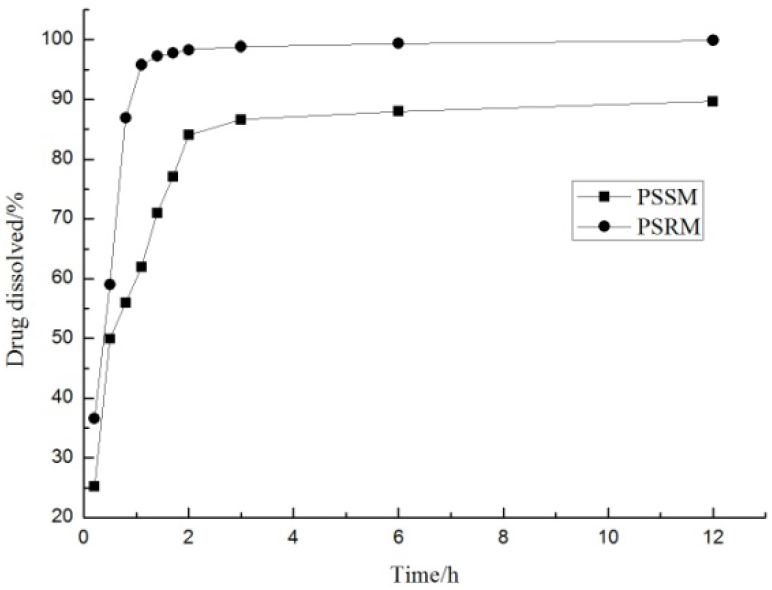
Release curve of PSSM and PSRM in

**Table 4.**
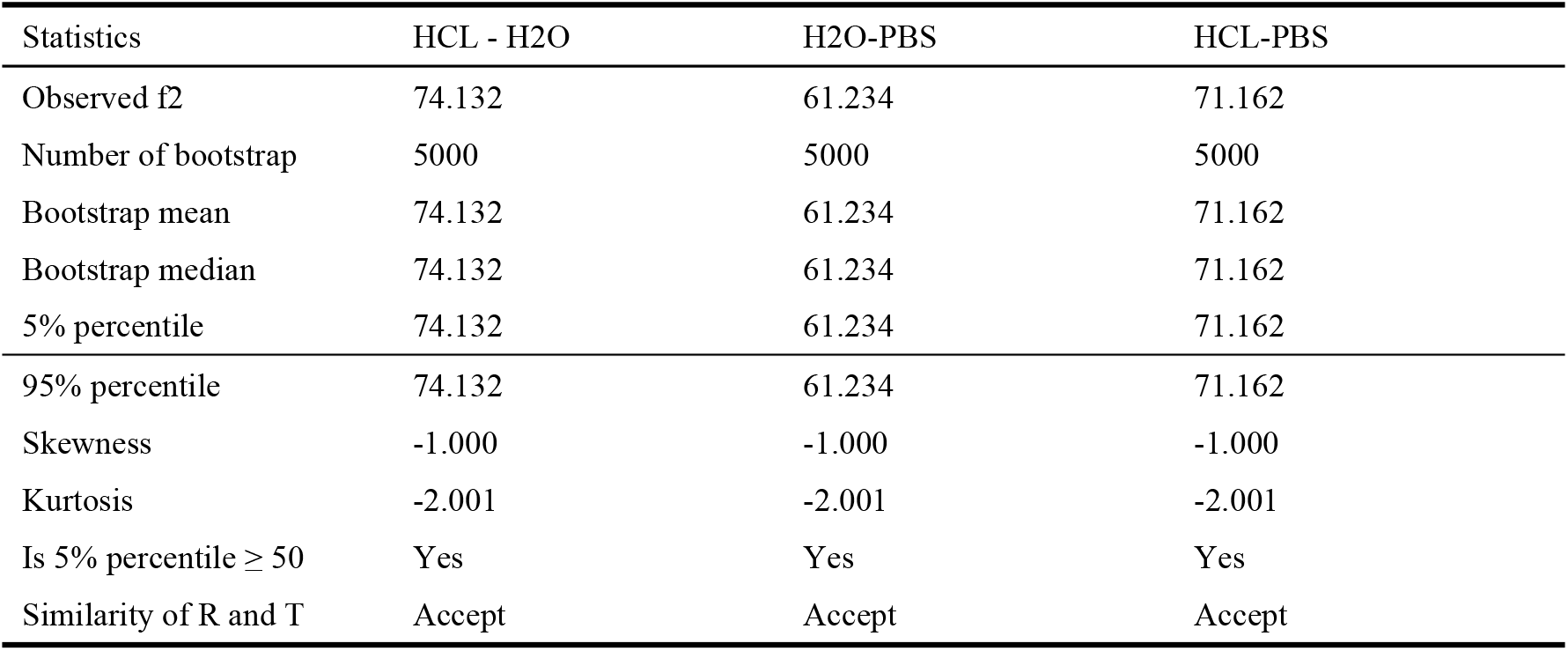
In-vitro Dissolution Profile Comparison in different diluents

| Statistics | HCL - H2O | H2O-PBS | HCL-PBS |
| --- | --- | --- | --- |
| Observed $f_2$ | 74.132 | 61.234 | 71.162 |
| Number of bootstrap | 5000 | 5000 | 5000 |
| Bootstrap mean | 74.132 | 61.234 | 71.162 |
| Bootstrap median | 74.132 | 61.234 | 71.162 |
| 5% percentile | 74.132 | 61.234 | 71.162 |
| 95% percentile | 74.132 | 61.234 | 71.162 |
| Skewness | -1.000 | -1.000 | -1.000 |
| Kurtosis | -2.001 | -2.001 | -2.001 |
| Is 5% percentile $\geq 50$ | Yes | Yes | Yes |
| Similarity of R and T | Accept | Accept | Accept |

**Table 5.** Regression equation and data fitting of PSSM release in pH 6.8 PBS

| Dissolution Data Modeling | Equation | Rsqr |
| --- | --- | --- |
| Zero-order with F0 | $F=58.524+3.628*t$ | 0.3973 |
| First-order with Fmax | $F=87.657*[1-\text{Exp}(-1.340*t)]$ | 0.9665 |
| Higuchi with F0 | $F=42.379+18.135*t^{0.5}$ | 0.6121 |
| Hixson-Crowell with Tlag | $F=100* [1-[1-0.098*(t+1.570)]^3]$ | 0.7686 |
| Korsmeyer-Peppas with F0 | $F=-71592.330+71653.858*t^{0.000}$ | 0.8695 |

#### Pharmacokinetic study

As shown in Fig. 13, after taking the same dose of PSSM and PSAM orally, the concentration of PS in PSSM group remained at 0.79-2.87 ug/ml within 24 hours, which was significantly higher than that in PSAM group (0.12-1.36 ug/ml). The Cmax value of PSSM group was 2.89±0.20 at Tmax=2.63±0.36 h, which was 1.55 times higher than that of PSAM group at Tmax=1.70±0.12 h(Table 6). The CL of PSSM group (3.94±0.87 ug/h) was significantly lower than that of PSAM group (15.35±1.41 ug/h). The bioavailability of PSSM group was 389.68% compared with that of PSAM group. Therefore, PSSM can significantly improve the bioavailability of PS.

**Fig 13.**
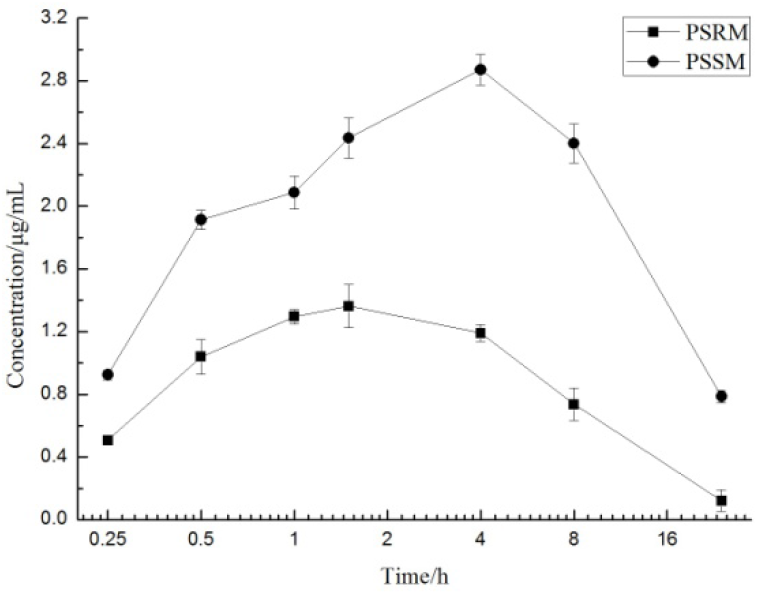
Mean plasma concentration-time profiles of PS in rats

**Table 6.** Pharmacokinetic parameters of PSSM and PSAM

| Formulation | AUC($\mu\text{g/mL}$)*h | Tmax(h) | Cmax($\mu\text{g/mL}$) | CL($\mu\text{g/h}$) | F(%) |
| --- | --- | --- | --- | --- | --- |
| PSSM | $59.66\pm12.24$ | $2.63\pm0.36$ | $2.89\pm0.20$ | $3.94\pm0.87$ | 389.68 |
| PSAM | $15.31\pm1.41$ | $1.70\pm0.12$ | $1.41\pm0.05$ | $15.35\pm1.41$ | |

## 4. Conclusion

In this study, by measuring the oil and water distribution coefficient of phytosterol, we know that phytosterol is not easily absorbed by human intestinal tract, so it is necessary to carry out studies on microemulsion preparation of phytosterol. In the process of formulation optimization of PSSM, solubility test, drawing of pseudo ternary phase diagram and particle size and potential test were carried out, and the preparation prescription of PSSM was determined as follows: lemon essential oil in oil phase, polyoxyethylene hydrogenated castor oil 40 and Tween 60 in emulsifier, polyethylene glycol 400 in co-emulsifier, Km = 7:3, Kp = 3:1, Ke = 50%. The pharmacokinetic analysis of PSSM synthesized by this method shows that PSSM can increase the bioavailability of PS more than three times, so it is necessary to do more in-depth research on the self-microemulsion delivery system of phytosterols.

## Abbreviations

LDL: : low density lipoprotein;
POP: : phytosterol oxidation product;
PS: : phytosterols;
SMEDDS: : self-microemulsifying drug delivery system;
PSSM: : phytosterol self-microemulsion;
LEO: : lemon essential oil;
HCO-40: : polyoxyethylene hydrogenated castor oil 40;
PEG400: : Polyethylene glycol 400;
IPM: : isopropyl myristate;
PEG400: : polyethylene glycol 400;
EL: : polyoxyethylene castor oil;
LA: : linoleic acid;
OA: : oleic acid;
EO: : ethyl oleate;
PSRM: : phytosterol raw materials;
HPLC: : high performance liquid chromatography;
DL: : drug loading;
EE: : entrapment efficiency;
P: : oil-water partition coefficient;
Km: : the ratio of HCO-40 toTween 60;
Kp: : the ratio of emulsifier to co-emulsifier;
Ke: : emulsifier content;
PBS: : phosphate buffer;
TEM: : transmission electron microscopy;
PDI: : Polydispersity index

## Acknowledgements

We sincerely thank the Yuhua Guo, Yun Xu, Xue Long, Jing Jin for technical assistance.

## Availability of data and materials

All data generated or analyzed during this study are included within the article.

## Authors’ contributions

Chuanxun Yuan and Xueru Zhang designed the study; Xueru Zhang, Risheng Jin and Peng Sun analyzed the biochemical data; Xueru Zhang and Peng Sun analyzed the Histopathological data. Chuanxun Yuan, Xueru Zhang, Risheng Jin drafted the manuscript. All authors read and approved the final manuscript.

## Ethics approval

All animals used in this study were treated strictly according to the National Institutes of Health Guidelines of the Care and Use of Laboratory Animals. All procedures were approved by the Animal Care Review Committee, Hefei University of technology, China.

## Consent for publication

Not applicable.

